# An optimised protocol for the relative quantification of ATP and polyphosphates in the microalga *Chlamydomonas reinhardtii*

**DOI:** 10.1101/2024.12.16.628749

**Authors:** Eliora Israelievitch, Alix Boulouis

## Abstract

**Background:** Polyphosphates (polyP) and ATP are phosphate-containing metabolites present in prokaryotes and eukaryotes. PolyP has a wide variety of functions including phosphate and cation storage. ATP is a central metabolite in cellular bioenergetics and the phosphate providing substrate of polyP. In the green microalga *Chlamydomonas reinhardtii*, polyP synthesis is suggested to buffer ATP concentration, and the role of polyP in energetic metabolism requires further investigation. In this aim, relative quantification of both metabolites is needed. Because ATP and polyP half-lives differ greatly, harvesting generates biases in this relative quantification in current methods. For this reason, we present here a joint protocol optimised to compromise between maximal yield and specific constraints of both assays.

**Methods:** The optimised method quantifies ATP and polyP from the same *C. reinhardtii* cell extract after neutral phenol-chloroform extraction. Cells are directly pipetted from the culture to the phenol-chloroform- EDTA extraction mix. After a second chloroform extraction, ATP is quantified directly from the extract, while polyP measurement requires purification by ethanol precipitation. We used one-way analysis of variance or Kruskall-Wallis testing and appropriate post hoc testing to evaluate statistical effects in our results.

**Results:** We show that he polyP/ATP ratio of the reference strain CC-4533 in exponential mixotrophic growth is around 65. While the optimised protocol performs as well as specific protocols for either ATP or polyP, the dispersion of the polyP/ATP ratio is twice better than for the separate metabolites. Direct sampling from the culture works better than centrifugation and filtration to maintain physiological conditions and high polyP yield. Using spiking with ATP and polyP, we show that ATP and longer chain polyP are fully recovered but not very short chain polyP. Finally, we show that the polyP chain length distribution extracted from CC-4533 is very broad, reaching up to several thousand P with a mean around 200 P.

**Discussion:** Our protocol improves the precision of relative quantification of ATP and polyP by using neutral phenol-chloroform extraction and allows polyP/ATP ratio calculation from low-density samples and without normalisation. It can be applied to other microorganisms or cells, in a variety of physiological and stress conditions.

## List of abbreviations

ATP: adenosine triphosphate
polyP: polyphosphates
TCA: trichloroacetic acid

## Introduction

Polyphosphates (polyP) are polymers of inorganic phosphate (Pi) believed to have pre-existed life on Earth (1). From phosphate and cation storage to cell signalling and stress response, polyP have a wide range of functions in cells (2,3). PolyP are found in all life kingdoms, but their synthesis differs between prokaryotes and eukaryotes (3). In eukaryotes, the vacuolar transporter chaperone enzyme (VTC) polymerises polyP from ATP (3). ATP also has a central role in cellular bioenergetics, and provides energy in numerous metabolic reactions, making ATP concentration critical to cell physiology. In the green microalga *Chlamydomonas reinhardtii*, a model species for bioenergetics, a mutant without the VTC enzyme accumulates higher levels of ATP, in conditions where the wild-type produces polyP (4). This led Sanz-Luque et al. to suggest that polyP synthesis allows to buffer the excess ATP in *C. reinhardtii* (4). Additionally, an indirect participation of polyP in energy storage in eukaryotic cells has been suggested (4,5). Testing the hypothesis that polyP regulate cell bioenergetics requires high accuracy in the quantification of the relative amounts of polyP and ATP in cells. In order to achieve this, the extraction method should be as similar for both metabolites as possible, in order to avoid biases introduced by the different steps of cell harvesting, extraction itself and purification. Stability of both metabolites should be taken into account at each step of the protocol, as well as assay-related constraints.

Several protocols already exist to extract and quantify either ATP or polyP, mainly consisting in their separation from the protein fraction. For ATP, the extraction is usually based on acids like trichloroacetic acid (TCA) or perchloric acid (6,7). However, Chida et al. showed on mammalian tissues and blood cells that a neutral phenol-chloroform based extraction is more efficient (8). For polyP, early extraction protocols (in the 1980’s and 1990’s) have been based on acids, mainly TCA, carbon tetrachloride or phenol-chloroform, although only the latter properly retrieved granular polyP (9,10). More recently, two groups optimised a polyP extraction method in *Saccharomyces cerevisiae* based on neutral phenol-chloroform to decrease the number of steps while increasing the efficiency (11,12). Thus, a neutral phenol-chloroform-based extraction method seems very suitable for both ATP and polyP. In this method, variability comes from the unprecise retrieving of the aqueous phase at each extraction step. This becomes irrelevant for the relative quantification of ATP and polyP, alleviating the need of a common normalisation. Indeed, normalisation usually uses cell count, chlorophyll concentration or dry mass, all parameters that can vary with cell cycle, light and nutrient conditions (13).

Biological material harvesting is a key step of metabolite extraction. This is especially true for microorganisms growing in liquid cultures, since the metabolite concentrations of the whole culture are largely diluted by the medium. ATP, a very short-lived metabolite in physiological conditions (14), requires a quasi-instantaneous harvesting and quenching method. Direct sampling of *C. reinhardtii* cells from their culture medium has been previously used, rapidly mixing the algal culture with very cold methanol (15). On the contrary to ATP, polyP is a more stable metabolite that can be extracted after steps of cell centrifugation and washing (12). This procedure ensures that only intracellular polyP is taken into account in the measurement and that Pi contained in the medium does not contaminate the Pi-based colorimetric assay of polyP after hydrolysis to Pi. Centrifugation and washing might or not affect *C. reinhardtii* polyP stocks, but will surely have an impact on cell physiology by changing light intensity, oxygen availability and medium composition, thus representing a potential threat to correct ATP quantification. Another option to eliminate medium and cell Pi contribution to the assay is to purify polyP from the cell culture extract by ethanol precipitation (11). In order to use this method, one should check that this additional step specific to polyP does not introduce a bias in the relative quantification of ATP and polyP.

We optimised a method to measure physiological concentrations of ATP and polyP in CC-4533 *C. reinhardtii* cells, largely inspired by Chida et al. (8) for the extraction part, and we compared it with other available protocols. We demonstrate the efficiency of the method for both metabolites. We studied the impact of the cell harvesting method on the final ATP and polyP quantifications and on cell physiology. We show that using a precipitation step after extraction allows to purify polyP from contaminating Pi without affecting total polyP quantification, thus allowing to directly harvest cells in their culture medium. Additional minor optimisation steps concerning EDTA concentration and sample storage are also described. The resulting joint extraction protocol for ATP and polyP thus offers the possibility to calculate polyP/ATP ratios independently of any normalisation method.

## Methods

### C. reinhardtii strain and cultivation

The CC-4533 (CMJ030) strain was cultivated mixotrophically in a medium composed of 1 mM phosphate, 17.5 mM glacial acetic acid, Beijerincks nutrient solution, 15 mM HEPES buffer and Kropat traces, adjusted to pH 7 with KOH, in 50 mL liquid cultures, under low constant illumination (∼10 µmol photons m^-2^ s^-1^) and constant agitation (130 rpm). Cultures were inoculated at 1-5 10^5^ cells mL^-1^ and let grow until log phase (0.5-3 10^6^ cells mL^-1^), where they were maintained at least two days by dilutions before each experiment. Cells used for experiments were in log phase at ∼2 10^6^ cells mL^-1^.

For data normalisation, cell density in the culture was measured before each experiment with a haemocytometer after killing cells with 5% iodine tincture. To take into account potential losses of material in the harvesting steps, cell density was quantified again after harvesting with optical density measurements at 550 nm, using the conversion factor of 8.3 10^6^ cells OD550^-1^ obtained by calibration of the measurement on CC-4533 (16).

### Protocols published for ATP extraction

All steps were performed on ice or at 4°C for centrifugation.

#### “Chida” method (8)

A volume of 300 µL of cell culture was added to 1.5 mL of pre-cooled extraction mix (phenol-chloroform-water 6:2:2, 12 µl of 50 mM EDTA) in a 2 mL tube. After 20 s of vortexing, the tube was briefly centrifuged and the lid was filled with silicone vacuum grease to easily separate the aqueous and organic phases. After centrifugation (12000 g, 5 min), the aqueous phase was transferred to a pre-weighed 1.5 mL tube. The tube was weighed again to measure the exact extract volume and kept at −20°C until polyP precipitation and ATP quantification.

#### “Promega” method

The following TCA based extraction was performed according to the protocol supplied by the manufacturer (Enliten® ATP Assay System, Promega, USA). A volume of 400 µL of cell culture was added to 400 µL of cold 10 % trichloroacetic acid (TCA). After 20 s of vortexing, the suspension was left on ice for ca. 1 hour. After another 20 s vortexing, 400 µL of TA buffer (1 M acetate, 10 mM Tris, pH 7.8) were added to the mix. The tube was centrifuged (10000 g, 5 min) and 20 µL of the supernatant were transferred in a new 2 mL tube containing 1980 µL of TA buffer. 10 µL of the suspension was deposited on pH paper to verify neutrality (pH should be approximately 7.8). The sample was kept at −20°C until polyP precipitation and ATP quantification.

### Protocols published for PolyP extraction

#### “Christ” method (12)

All steps were performed at room temperature (RT).

Cells were harvested by centrifuging (5000 g, 10 min) 1 mL of culture in a 2 mL tube and washed twice in milliQ water. The pellet was resuspended in 500 µL ME buffer (25 mM MOPS, 2.5 mM EDTA, pH 7) and 100 µL were used for optical density measurement. 300 µL of phenol were added to the remaining 400 µL of suspension. After 20 s vortexing, the tube was incubated 10 min at 45°C, cooled for 2 min on ice and briefly centrifuged. 1 mL of chloroform was added and vortexed 20 s to extract the phenol dissolved in the aqueous phase. After a brief centrifugation, the lid was filled with silicone vacuum grease. After centrifugation (12000 g, 5 min), the aqueous phase was transferred to a new pre-weighed 1.5 mL tube. The tube was weighed again to measure the exact extract volume and kept at −20°C until polyP precipitation and ATP quantification.

#### “Bru” method (11)

Cells were harvested by centrifuging (5000 g, 5 min, 4°C) 1 mL of culture in a 2 mL tube. The pellet was resuspended in 500 µL AE buffer (50 mM sodium acetate pH 5.3, 10 mM EDTA) and 100 µL were used for optical density measurement. Volumes of 300 µL of phenol and 40 µL of 10 % SDS were added to the suspension. The tube was mixed (4 inversions and 5 s vortexing) and incubated 5 min at 65°C, then cooled on ice for 1 min. A volume of 300 µL of chloroform was added before mixing again. The tube was briefly centrifuged and the lid was filled with silicone vacuum grease. After centrifugation (13000 g, 2 min, RT), the aqueous phase was transferred to a new 1.5 mL tube containing 500 µL of chloroform and mixed, then briefly centrifuged. The lid was filled with silicone vacuum grease. After centrifugation (13000 g, 2 min, RT), the aqueous phase was transferred to a new pre-weighed 1.5 mL tube. The tube was weighed again to measure the exact extract volume. 3 µL of RNAse A (10 mg mL^-1^) and 3 µL of DNAse I (10 mg mL^-1^) were added, incubated 1 h at 37°C and kept at −20°C until polyP precipitation and ATP quantification.

### Optimised extraction protocol

All steps were performed on ice or at 4°C for centrifugation.

A volume of 600 µL of cell culture was added to 1.2 mL of pre-cooled extraction mix (phenol- chloroform 3:1, 12 µL of 50 mM EDTA) in a 2 mL tube. After 20 s vortexing, the tube was briefly centrifuged and the lid was filled with silicone vacuum grease. After centrifugation (12000 g, 5 min), the aqueous phase was transferred to a new 2 mL tube. A volume of 1 mL of chloroform was added and vortexed 20 s to extract the phenol dissolved in the aqueous phase. The tube was briefly centrifuged and the lid was filled with silicone vacuum grease. After centrifugation (12000 g, 5 min), the aqueous phase was transferred to a new pre-weighed 1.5 mL tube. The tube was weighed again to measure the exact extract volume and kept at −20°C until polyP precipitation and ATP quantification.

### Cell harvesting using filtration

A volume of 1 mL of cell culture was filtered through a 3 µm mesh filter (Nuclepore Track-Etched Membranes, Whatman, USA) using a 15 mL glass filtration unit (Schott Duran, Germany). A 50 mL syringe was used to slowly extract air from the lower compartment of the filtration unit, in order to stress cells the least possible. If the sample was washed, cells were resuspended in 1 mL of milliQ water and filtered again. The filter was then inserted in a 2 mL tube and cells were harvested with 750 µL ME buffer (MOPS 25 mM, EDTA 2.5 mM, pH 7).

### Photosynthesis measurement

The maximal photosystem II (PSII) yield was measured with a Speedzen fluorescence imaging setup (JBeamBio/API, France). A volume of 50 µL of cell suspension was put on a metallic plate and inserted in the Speedzen. The plate was left in the dark for 2 min prior to measurement, which consisted of minimal fluorescence (*F*_0_) measurement in the dark and maximal fluorescence measurement (*F*_m_) after a saturating pulse (2600 µmol photons m^-^² s^-1^, 250 ms of green LEDs at 532 nm). The maximal PSII yield (Fv/Fm) was calculated using the formula of Eq. 1:

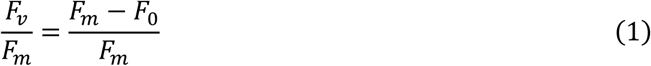

### PolyP precipitation

A volume close to 300 µL of extract, corresponding to 0.5-1 10^6^ cells, was used for polyP purification by ethanol precipitation. The precise volume of extract for precipitation was calculated for each sample to account for potential cell amount differences, in order to fall into the detection range of the polyP concentration assay. The extract was added to a pre-cooled 2 mL tube containing 2.5 volumes of cold 100 % ethanol and 0.1 volume of 3M sodium acetate (pH 5.3), and precipitation occurred overnight at −20°C, unless stated otherwise. After washing in 70% ethanol, the pellet was dried at room temperature for 10 min, then resuspended in 220 µL of milliQ water.

### Assays

#### ATP bioluminescent assay

ATP concentration in extracts was measured with the firefly Luciferin-Luciferase assay kit (Enliten ATP Assay System, Promega, USA). Extracts were diluted 100 times in ATP-free water. A volume of 100 µL of diluted extracts and pure ATP standards was pipetted onto a dark flat-bottom 96-well plate. Just before measurement, 100 µL of reagents were added to the measured well and mixed twice with the pipet. Measurements were performed in a CLARIOstar microplate reader (BMG Labtech, Germany). Luminescence at 555-70 nm was measured for 10 s to maximise signal detection.

#### PolyP colorimetric and fluorometric assays for concentration and average chain length

PolyP concentration in extracts was measured by enzymatic hydrolysis with scPpx1p and scIpp1p and colorimetric Pi detection, using the Phosfinity Total Polyphosphate Quantification Kit (Aminoverse, Netherlands) as described in (17). Briefly, 100 µL of purified polyP and positive control were pipetted in two separate wells of a transparent flat-bottomed 96-well plate, as well as 100 µL of Pi standards and negative control. A volume of 50 µL of enzyme buffer was added to only one of the two wells of each sample and positive control. Then 50 µL of a premix of enzyme buffer and enzymes (49:1) were added to all the other wells. After mixing two times with a multipet, the plate was incubated 1 h at room temperature. A volume of 50 µL of detection solution (based on molybdate and ascorbate) was distributed to all wells and mixed twice with a multipet. The plate was inserted in the CLARIOstar microplate reader (BMG Labtech, Germany) and incubated 2 min at room temperature before reading absorbance at 882 nm.

PolyP average chain length was determined by hydrolysis of polyP down to polyP2 with scPpx1p, conversion to ATP with ATP sulfurylase, then to glucose with hexokinase then to NADPH with glucose 6-phosphate dehydrogenase, and subsequent NADPH fluorometric detection, using the ChainQuant Polyphosphate Chain Length Determination Kit (Aminoverse, Netherlands) as described in (17).

Briefly, 100 µL of purified polyP were pipetted in two separate wells of a dark flat-bottomed 96-well plate, as well as 100 µL of polyP2 standards and positive and negative controls in one well. 50 µL of a premix of enzyme buffer, additive and enzymes (44:5:1) without scPpx1p and ATP sulfurylase were added to only one of the two wells of each sample. A volume of 50 µL of a premix of enzyme buffer, additive and enzymes (44:5:1 or 42:5:3 for cell extracts) was added to all the other wells. Note that we only obtained consistent chain length results in our cell extracts when we tripled the enzyme amounts, probably because of the rather long (up to several thousands P) polyP chains in our samples. After mixing two times with a multipet, the plate was incubated 1 h at room temperature and 5 min in the microplate reader (CLARIOstar). Finally, the fluorescence was measured (excitation at 340-9 nm, emission at 460-9 nm).

#### Polyacrylamide gel electrophoresis of polyP extracts

Denaturing PAGE analysis of polyP extracts was carried out like in (18) with some modifications. PolyP standards (0.2 µg of P14, P60, P130 and P700) and three quantities (1 µg, 0.2 µg and 0.1 µg) of purified cell extract were loaded on a 4-18 % polyacrylamide 8 M urea denaturing gel (13 cm height). As a qualitative reference, 1 µg of 1 kb+ DNA ladder (N0550, New England Biolabs, USA) was also loaded. The gel was run overnight at 20 V and 3 h at 100 V. After a fixation step (30 min under mild agitation in the fixation buffer composed of 50 mM Tris-HCl pH8, 25 % methanol and 5 % glycerol), the gel was stained in a bath containing 2 µg mL^-1^ DAPI in the fixation buffer for 30 min, and destained in fixation buffer for 45 min. The gel was placed in a Gel Doc XR+ Molecular Imager (Bio-Rad, USA) and UV illuminated until total bleaching of polyP-bound DAPI (10 min).

### Chemicals

All chemicals used were of analytical grade and all dilutions were made with high purity deionized water (18 MΩ cm resistivity) obtained from a milliQ water purification system (Millipore, MA). Phenol was water saturated and equilibrated to pH 8 (Aquaphenol, MP Biomedicals, USA).

Chloroform was a mixture of chloroform and isoamyl alcohol (24:1) (BioUltra, Supelco, Sigma-Aldrich, USA). The DNAse I and RNAse A enzymes and DAPI were purchased from Thermo Fisher Scientific (USA). Pure ATP standards for spiking were prepared from ATP standard of an ATP Colorimetric/Fluorometric Assay Kit (MAK190, Sigma-Aldrich, USA). Pure polyP standards for spiking were kindly given by RegeneTiss Inc. (P14, P60 and P130) or purchased from Sigma-Aldrich (P3) or Kerafast (P700). Silicone vacuum grease (High Vacuum Grease, Molykote, DuPont, USA) and all other chemicals were purchased from Sigma-Aldrich (USA).

### Data analysis and statistics

Data were treated by R scripts in RStudio. Statistical analysis included ANOVA and Tukey HSD tests for data with homogeneous variances (Fig. 1A, 2-4) and Kruskal-Wallis and Dunn tests for data with non-homogeneous variances (Fig. 1B and 5). Variance homogeneity was tested with a Fisher’s test.

**Figure 1:**
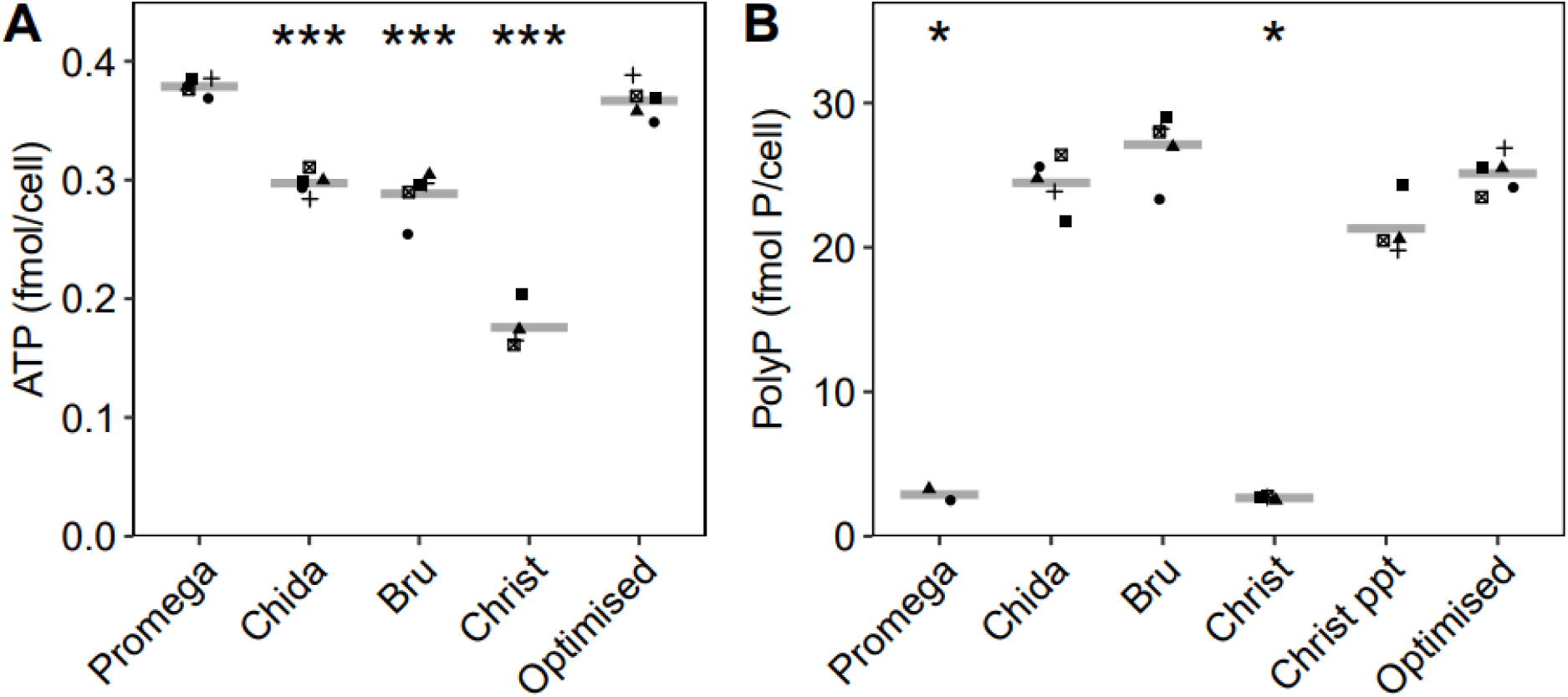
Cellular concentrations of ATP and polyP after extractions with previously published protocols and the optimised protocol. ATP (A) and polyP (B) were assayed after metabolite extraction using phenol-chloroform (“Bru” (11), “Chida” (8), “Christ” (12) and optimised protocols) or TCA-based (“Promega”) protocols, from the same CC-4533 culture. Four to five biological replicates of the same initial culture were measured. Three out of five replicates of the Promega protocol had non measurable amounts of polyP. Grey crossbars indicate the mean values. “ppt” stands for precipitation of polyP after the Christ protocol. Stars represent significant differences compared to the optimised protocol (one-way ANOVA for ATP and Kruskal-Wallis test for polyP, corrected p-value coding: * p < 0.05, ** p < 0.01, *** p < 0.001).

**Figure 2:**
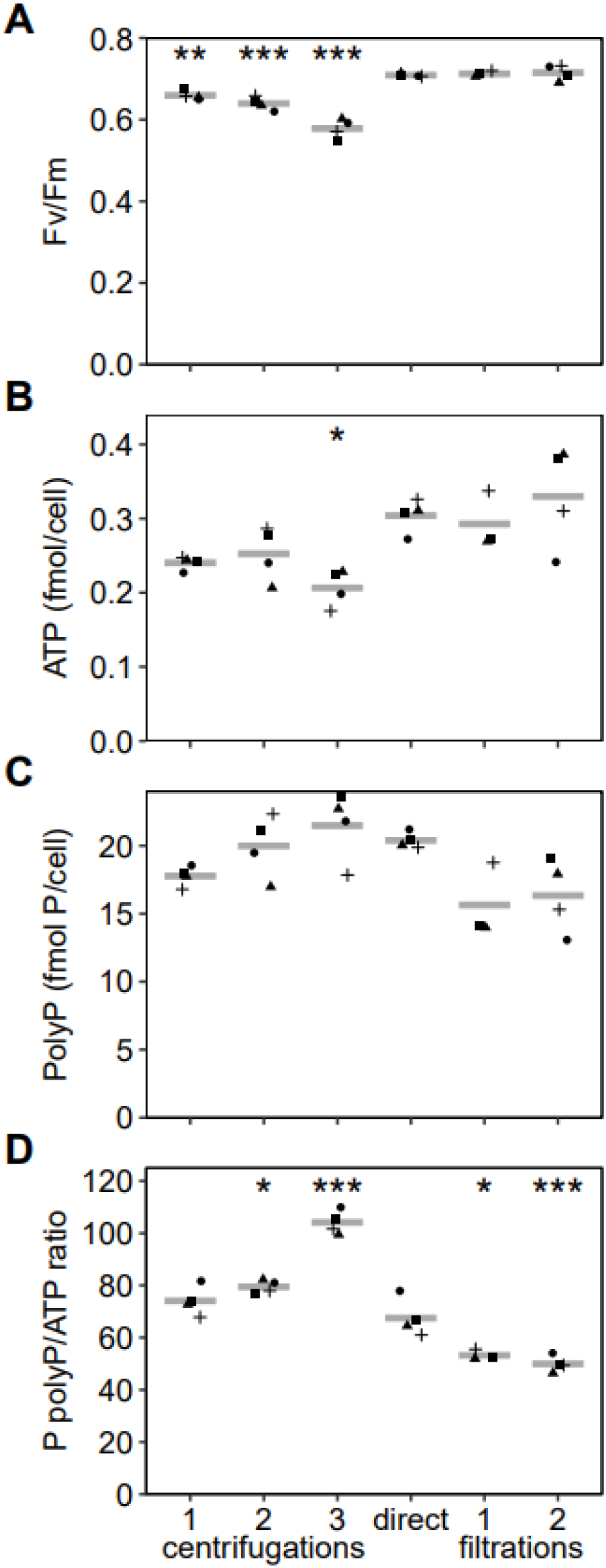
Comparison of harvesting methods for ATP and polyP measurements. Cells were harvested by one or several rounds of centrifugation or slow filtration, or directly from the culture. Cells were washed with milliQ water between harvesting steps in the case of several centrifugations or filtrations, and ME buffer was used for the last resuspension of cells. After harvesting, fluorescence was measured to calculate the Fv/Fm ratio (A) and extraction with the optimised protocol was carried out to measure ATP (B), polyP (C) and calculate the polyP/ATP ratio (D). For photosynthesis measurement, samples of the “direct” protocol contained only culture solution, without EDTA. Four harvesting replicates (or three, for the 1 filtration protocol) of the same initial culture are shown. Grey crossbars indicate the mean values. Stars represent significant differences compared to direct harvesting (ANOVA: * p < 0.05, ** p < 0.01, *** p < 0.001).

**Figure 3:**
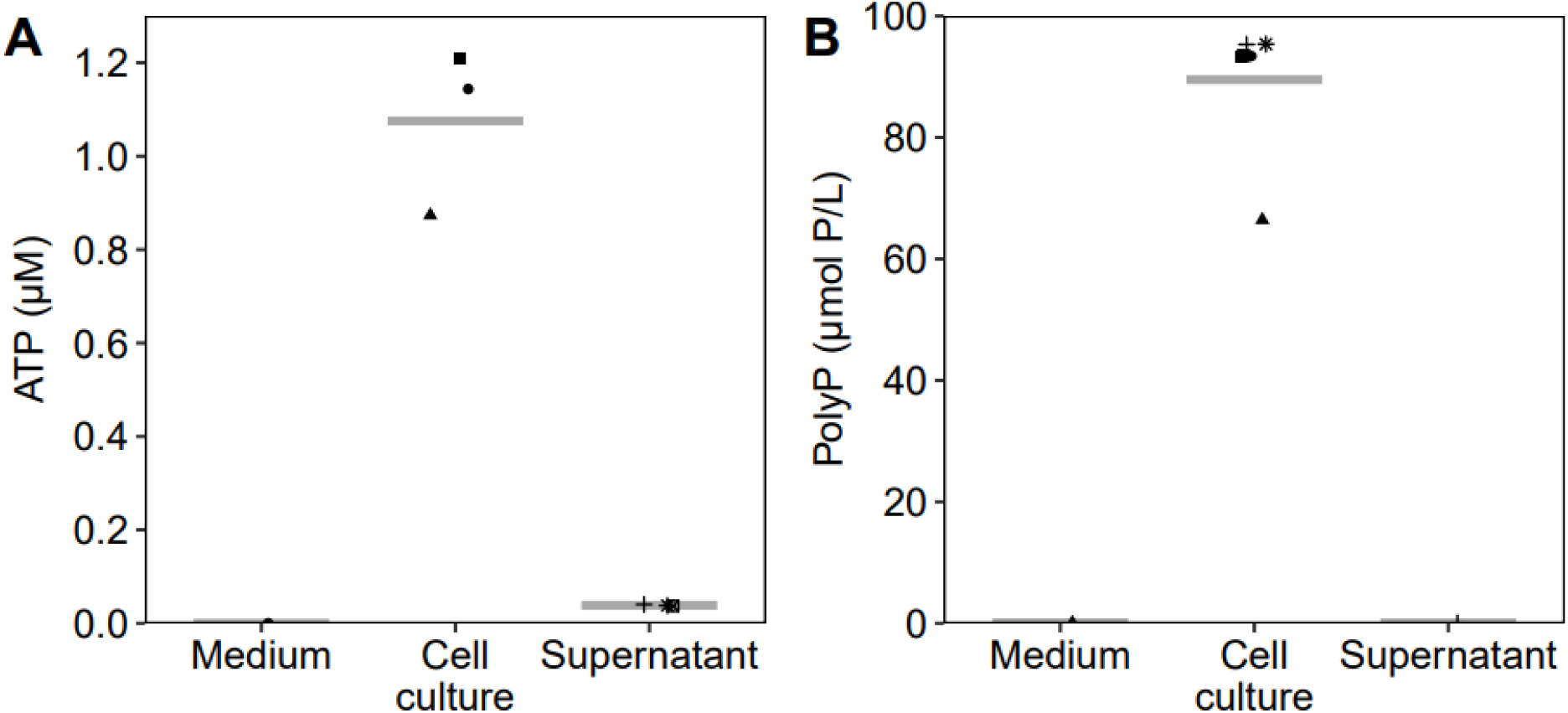
PolyP and ATP are not found in the extracellular medium of a *C. reinhardtii* culture. ATP (A) and polyP (B) were extracted from equal volumes of fresh new medium, a log-phase cell culture and the supernatant of this culture after removing cells by centrifugation (5 min, 12000 rcf). Three replicates for fresh medium and three to six replicates for cell and supernatant parts of the same culture are shown. Grey crossbars indicate mean values.

**Figure 4:**
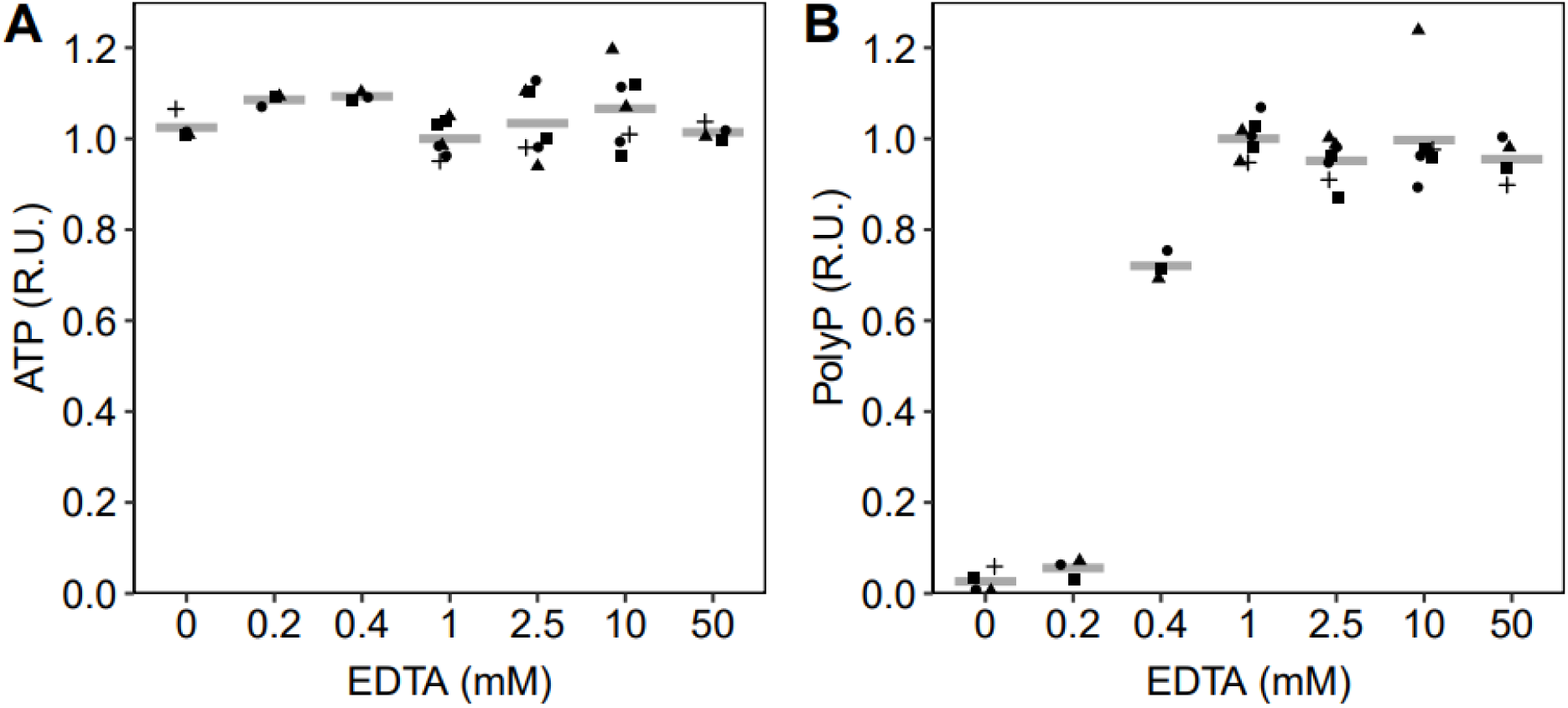
Dependence on EDTA concentration of ATP and polyP extraction efficiency. ATP (A) and polyP (B) were extracted with the optimised protocol with different EDTA concentrations. Data from two independent experiments with respectively three and four biological replicates were normalised to the mean value at 1 mM EDTA. Grey crossbars indicate mean values.

**Figure 5:**
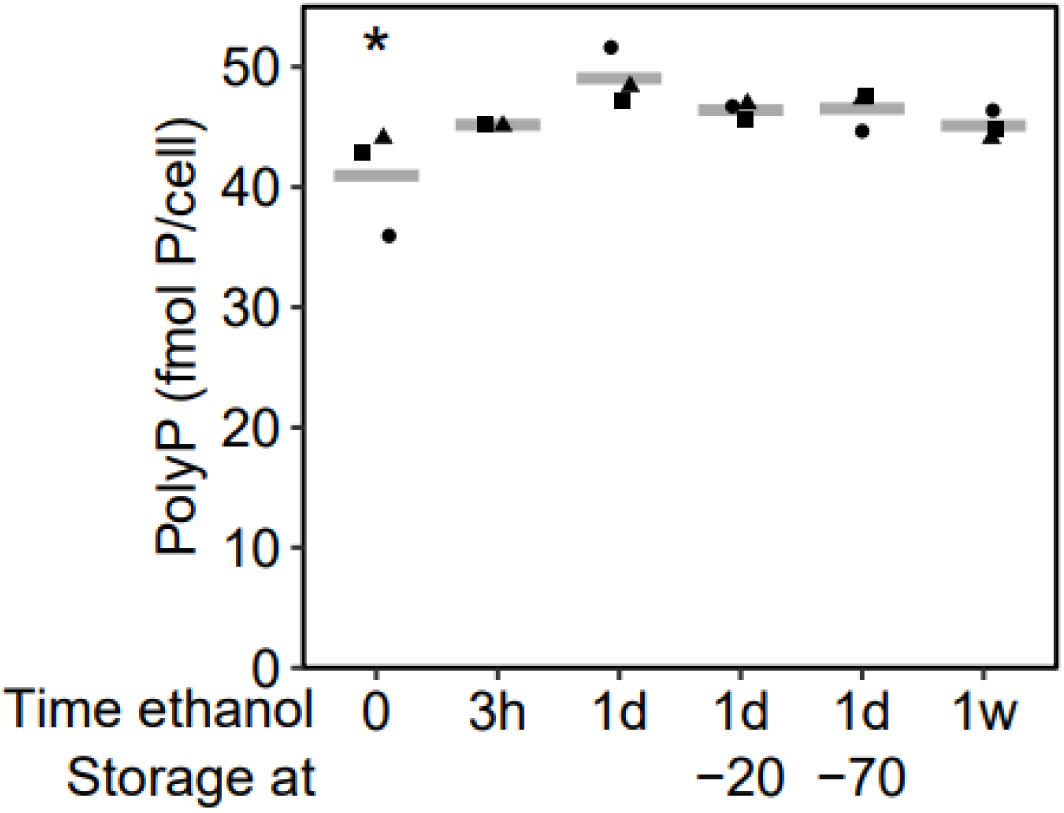
Comparison of different methods for conservation post extraction. Cells were harvested and washed twice with milliQ water before extraction to avoid P contamination of the assay. For protocol “direct”, polyP were assayed directly after extraction. For all the other protocols polyP were precipitated in ethanol, for three hours (3 h), overnight (1 d) or over one week (1 w). After the overnight precipitation, part of the purified extracts resuspended in water were stored at −20°C or −70°C for one week. Three biological replicates are shown. Grey crossbars indicate the mean values. The star represents a significant difference compared to overnight precipitation (ANOVA, * p<0.05).

Retrieved metabolite amounts in recovery experiments were calculated differently depending on the presence of cells in the medium. For ATP and polyP recovery rates from spiked fresh medium, we used Eq. 2:

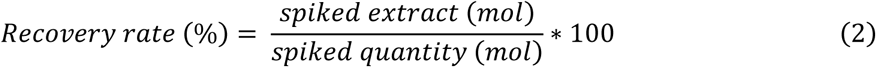

ATP and polyP recovery rates from the spiked culture were calculated for each pair of spiked replicate i and unspiked replicate j, according to Eq. 3:

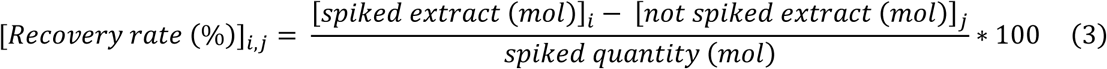

## Results

### Comparison of different ATP and polyP extraction methods

To develop a joint ATP and polyP extraction method for *C. reinhardtii*, we compared different methods published for ATP or polyP extraction with our optimised one, designated as “optimised” protocol in the rest of the study (Fig. 1). ATP extraction protocols were either based on phenol-chloroform (“Chida” protocol, (8)) or TCA (“Promega” protocol, also published for *C. reinhardtii* in (7)), while polyP extraction protocols were all phenol-chloroform based (“Bru” protocol (11) and “Christ” protocol (12)). We considered that the best protocols are those retrieving the highest amounts of each metabolite.

The optimised protocol performs as well as the best published protocols for each metabolite. For ATP extraction, the optimised protocol yields the same amount of ATP as the “Promega” protocol, whereas the “Chida”, “Christ” and “Bru” protocols extract significantly less ATP than the optimised protocol (Fig. 1A). For polyP extraction, the “Christ” and “Promega” protocols perform significantly worse than the others, retrieving hardly any polyP (Fig. 1B). The “Christ” protocol is originally a polyP extraction protocol, but it does not include a precipitation step. As we extracted 10 to 100 times less cells than in the publication and thus did not have to dilute samples before quantification, we also purified extracts from the “Christ” protocol before quantification (referred as “Christ ppt” in Fig. 1B). Adding this step allowed the “Christ” protocol to perform as well as the optimised, “Chida” and “Bru” protocols, showing that starting the extraction on high cell density samples is key to spare oneself the polyP precipitation step.

### Optimisation of cell harvesting

We compared three different harvesting approaches, with variants (Fig. 2). We harvested cells by (i) centrifugation with zero to two intermediate milliQ water washes, (ii) filtration with zero or one intermediate milliQ water wash and (iii) direct pipetting from the culture. To assess the potential stress caused by the different methods, we measured the PSII maximal yield (Fv/Fm) of the samples as a proxy of their physiological state, just before addition of the extraction mix. Indeed, this parameter will lower if cells experience hypoxia during the harvesting, which might occur in darkness or at high cell density (19).

Pipetting cell suspension directly from the culture to the extraction mix is the best harvesting protocol for cell physiology, ATP and polyP preservation (Fig. 2A-C). Indeed, this protocol intrinsically allows to extract cells in the same physiological state as in the culture. We thus further compare the centrifugation and filtration protocols to this direct harvesting. Filtration performs well, preserving the Fv/Fm ratio and retrieving the same amount of ATP as the direct protocol (Fig. 2A and B). The polyP content after filtration is slightly decreased, although non-significantly, compared to the direct harvesting (Fig. 2C). Centrifugation is the least reliable technique to maintain ATP levels of *C. reinhardtii* cells, even performed at 4°C. One centrifugation is sufficient to decrease the Fv/Fm ratio (Fig. 2A), and the more centrifugations and washes, the larger the Fv/Fm decrease. The protocol with three centrifugations triggers a significant ATP decrease with 32 % less ATP than with direct harvesting (Fig. 2B). The one and two centrifugation protocols also show a ∼20 % decrease in ATP compared to direct harvesting, although this is not statistically significant. On the contrary, centrifugation does not affect the polyP stocks (Fig. 2C), yielding comparable polyP concentrations to the direct harvesting protocol, whichever the number of centrifugations.

The physiological polyP/ATP ratio of CC-4533 cells cultivated in mixotrophic conditions and at log- phase culture stage is around 65 (Fig. 2D). Filtrations decrease this ratio, due to the loss of polyP they trigger. On the contrary, centrifugations increase the polyP/ATP ratio, due to the decrease in ATP they induce. The advantage of the joint extraction of ATP and polyP is obvious: while average coefficients of variation over all harvesting methods are 12% for ATP and 11% for polyP, the one of the ratio is twice lower at 6%.

We next wondered if ATP or polyP are present as free metabolites in the culture medium, which would cause them to be harvested with the direct protocol. Indeed, extracellular polyP is present in cell wall during cytokinesis in *C. reinhardtii* (20), and could end up in the medium. We thus checked the potential presence of ATP and polyP in culture supernatant after centrifugation. No extracellular polyP and very low levels of extracellular ATP were retrieved, and they were both absent from fresh medium (Fig. 3).

### Optimisation of the simultaneous ATP and polyP extraction method

To further improve the optimised protocol, we verified the effect of EDTA concentration on both ATP and polyP recovery from *C. reinhardtii*. We also investigated the conservation and precipitation conditions for polyP quantification.

EDTA is required for extraction of polyP, but does not affect ATP extraction (Fig. 4). Any concentration of EDTA above 1 mM is suitable for polyP extraction (Fig. 4, Panel B). The limit concentration of EDTA concentration is close to 0.4 mM, where ca. 30 % less polyP is extracted than at 1 mM. Thus, 1 mM EDTA was chosen as the final concentration for the rest of this study. The EDTA concentration might be increased to ensure stability of samples with higher polyP contents.

Precipitation is required to retrieve the highest amount of polyP in our conditions (Fig. 5). To determine the effect of polyP precipitation, we harvested cells by centrifugation and washed them twice before extraction, as described in the “Christ” method. Samples that were not precipitated show a slightly lower amount of polyP compared to precipitation in ethanol. The time in ethanol or the freezer temperature of conservation after resuspension in water have no effect on the measured amounts of polyP. For practical reasons, extracts were precipitated during one day to one week and stored at −20°C for the rest of the study.

### Recovery of polyP and ATP after extraction

Using the optimised protocol, polyP and ATP recovery rates were tested in different conditions (Fig. 6). Three amounts (0.2 nmol, 0.5 nmol and 1 nmol ATP) of the ATP standard and two amounts (10 nmol and 50 nmol Pi as polyP) of five polyP standards (P3, P14, P60, P700, P130) were spiked in tubes already containing the extraction mix and either fresh medium or cell suspension. These spiking amounts were chosen to be close to or smaller than the amounts extractable from the cells they were mixed with. We tested conditions with and without cells, keeping in mind that these reconstitution systems with standards in solution introduced in medium probably only partially mimic the biochemical environment and the structure of endogenous polyP and ATP during extraction. Please note that in Fig. 6B and 6D, we display the polyP standards in the order of their mean sizes measured with the chain length assay (Fig. 7).

**Figure 6:**
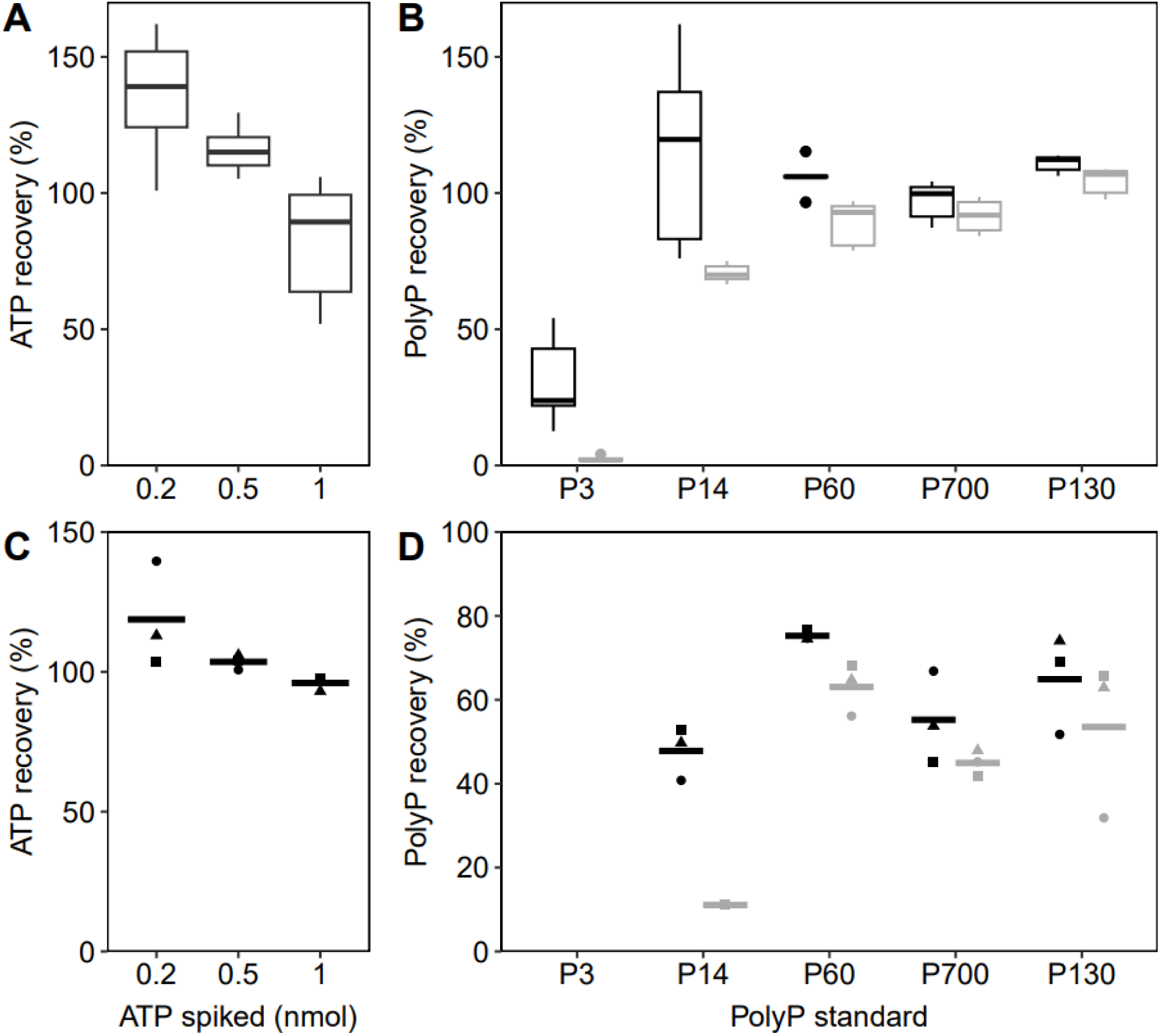
ATP and polyP recovery after spiking before extraction. Extractions of pure metabolites were done with (A, B) or without (C, D) CC-4533 cells. Three different quantities were introduced for ATP recovery (A, C) as shown on the x axis. Two different quantities were introduced for polyP recovery (B, D): 10 nmol P (black) and 50 nmol P (grey) alongside with five different polyP chain lengths as shown on the x axis. For extractions without cells, three biological replicates are shown and grey crossbars indicate mean values. For extractions with cells, boxplots represent the nine recovery rates calculated from cell extractions with and without spiking with three biological replicates each. Note that polyP standards are ordered according to their mean chain lengths and not their names.

**Figure 7:**
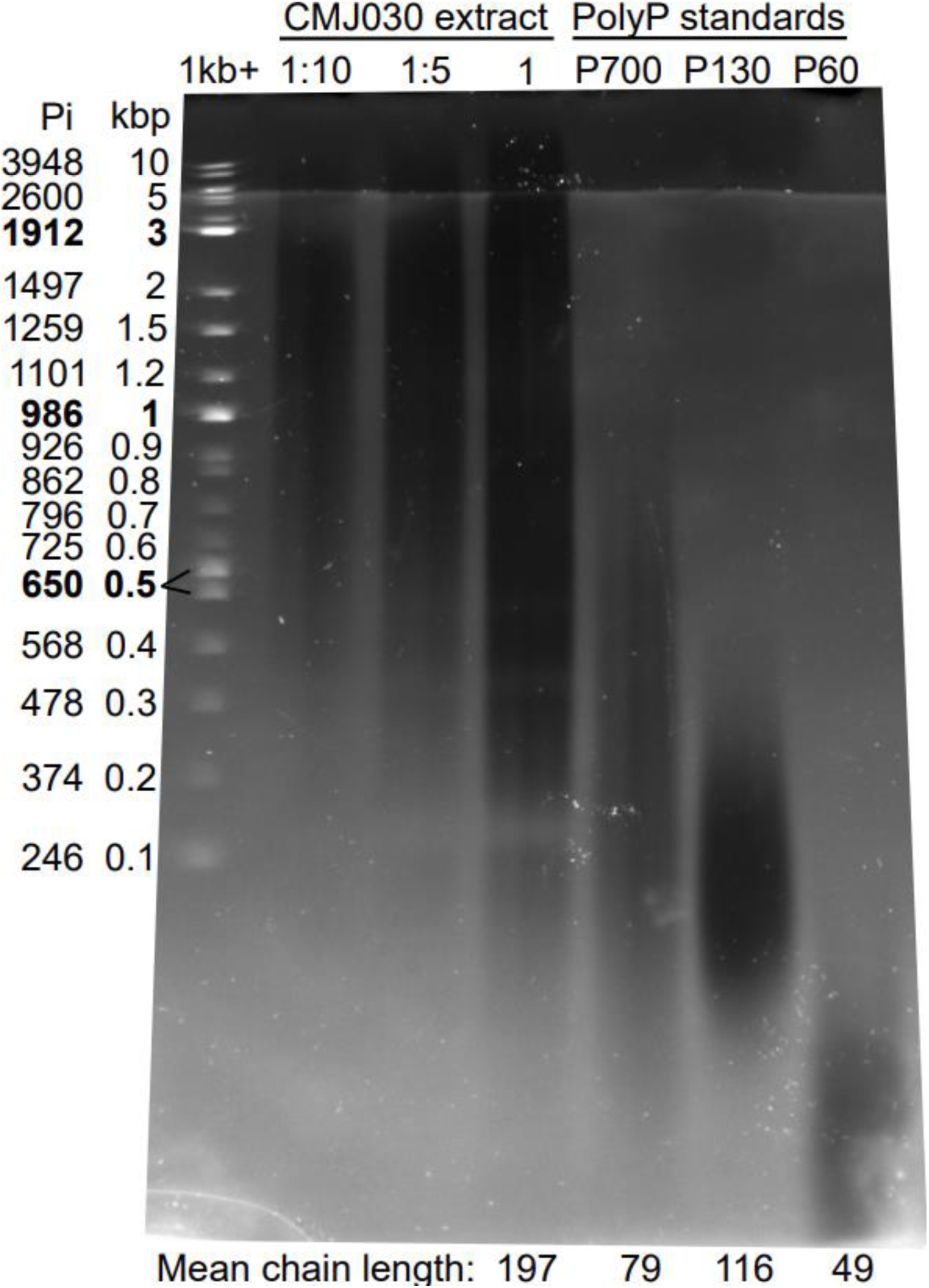
Polyacrylamide gel of CC-4533 cell extract and polyP standards. Cell extract of twelve extraction replicates from the same culture were pooled, from which three quantities (1 µg, 0.2 µg and 0.1 µg) were loaded on a 4-18% polyacrylamide denaturing gel. A mass of 0.2 µg of three polyP standards and 1 µg of the 1 kb+ DNA ladder (NEB) were also loaded for comparison. The depicted equivalent polyP chain lengths in P were calculated according to (21). Mean chain lengths displayed at the bottom of the gel are the results of the chain length enzyme assay.

ATP is fully recovered after the extraction with (Fig. 6A) or without cells (Fig. 6C). There is no effect of the presence of cells on ATP recovery during the extraction. Smaller ATP quantities are slightly overestimated in both conditions, especially in the cell-containing experiment.

PolyP with longer chains are fully recovered in cell-containing environment but not smaller chain polyP (Fig. 6B and D). Triphosphate (P3 in the figure) is not recovered at all (not measurable) when extracted in fresh medium (Fig. 6D) but partially (less than 50 %) recovered when extracted with cells (Fig. 6C). From P14 (for smaller spiked amount) or P60 (for larger spiked amount), the retrieved polyP quantity is maximal, reaching 100% in the presence of cells and only ∼70% without cells. Similarly to ATP, polyP recovery rates are smaller with higher spiked polyP amounts, regardless of polyP sizes.

Overall, polyP recovery rates are higher when extracted with cells than without.

### Chain lengths of polyP extracted from CC-4533

Given the apparent dependency to long chain of our extraction procedure, we then wondered how large the extracted polyP of *C. reinhardtii* are. We ran polyP extracted from CC-4533 and commercial polyP standards on denaturing PAGE (Fig. 7). We also measured the mean chain length on the same precipitated extract and on standards with an enzyme assay (17).

The CC-4533 strain of *C. reinhardtii* produces a large range of sizes of polyP, spanning and sticking out of the zone covered by the 1 kbp+ DNA ladder, with the maximal signal between 1000 and 2000 Pi residues. This is larger than commercially available polyP standard P700, which contains the largest polyP chains. Signal intensity on the gel being related to mass, this technique is not very well suited to evaluate the mean size of a polyP population. We thus quantified the mean chain lengths of the commercial standards and the CC-4533 extract with an enzyme assay (17). P3, P14, P60 and P130 standards have mean chain lengths similar to their names, with respectively 3 P, 16 P, 49 P and 116 P. The P700 standard has a mean chain length of 79 P, much smaller than its label, which is consistent with the wide size distribution observed on the gel. PolyP extracted from CC-4533 have a mean chain length around 200 P.

## Discussion

We optimised a protocol to simultaneously extract polyP and ATP directly from *C. reinhardtii* cells in culture. The method is simple to implement with few steps, and its efficiency compares with the published methods for polyP extraction optimised in other microeukaryotes (11,12,22). We identify some key parameters ensuring the efficiency of the method.

Extracts need to be clean of any phenol traces for both ATP and polyP quantification assays to work properly. PolyP purification by ethanol precipitation is paramount to properly quantify extracted polyP. This is especially true if the concentration in extracted samples is too low to necessitate further dilution for the assay, as performed in (12). In the case of lowly concentrated cultures (lag or exponential phase), precipitation allows both to get rid of phenol traces that could inhibit the assay and to concentrate the extracts. Indeed, the “Christ” extraction protocol is less efficient for samples lowly (e.g. five-fold) or not diluted (see Fig. 1 & 2 and (22)). Precipitation does not decrease the measured amount of polyP (Fig. 5), rather the opposite. This probably reflects the phenol removal effect, which would mask any loss of material during precipitation. Nevertheless, if any material is lost during precipitation, this is only very small chain polyP (Fig. 6), which would not bias very much the total P quantification, given the large size of *C. reinhardtii*’s polyP (Fig. 7 and (23)). However, we recommend to measure polyP size distribution of samples if the optimised method would be used with a different organism (e.g. (24) shows that photosynthetic microorganisms have various lengths of polyP, *Phaeodactylum tricornutum* having especially short ones). The optimised protocol extracts significantly more ATP than the original “Chida” protocol (Fig. 1). The main difference between the two protocols is the additional chloroform extraction step, which allows to decontaminate the aqueous phase from phenol residuals (25).

PolyP purification allows to concentrate the extracts for the quantification assay, making the optimised protocol usable on samples containing low polyP levels. Compared to the “Chida” protocol, the optimised protocol replaces water by cell suspension, allowing larger sample volumes to be extracted. This difference doesn’t affect the extraction efficiency (see Fig. 1) and allows to double the number of extracted cells. Thus, it is possible to follow cell polyP content during a growth curve with our method.

Slowly filtrating low-density culture cells, washing them with water and resuspending them in the presence of 1 mM EDTA do not affect cell physiology, as measured by the PSII maximal yield (filtration and direct protocols, Fig. 2). PSII maximal yield is affected by our centrifugation protocol, where we centrifuge cells at 4°C in order to slow down reactions with ATP. During centrifugation, cells are in the dark for several minutes while they fall into hypoxia, becoming unable to produce ATP by mitochondrial respiration for a period of time three orders of magnitude above its life-time in the cell (14). This and plain shear stress could explain the loss of ATP and the decrease of Fv/Fm after such treatment. Another possible stress source is the cold temperature we used in the centrifuge to slow down enzymatic reactions. Although short-term effects of cold stress have not yet been sufficiently studied, cells acclimated for one day at 4°C have a two-fold reduced Fv/Fm and a more than two-fold starch increase (26,27).

Our protocol retrieves less metabolites when the spiked quantity is larger (Fig. 6). For both ATP and polyP, we introduced amounts similar to expected amounts in the co-extracted cells, and smaller amounts. It is hard to imagine that doubling the metabolite quantity to be extracted saturates the extractability capacity. This intriguing effect was already observed with a different method of extraction, without an explanation (28). Whereas the presence of cells does not affect the recovery of spiked ATP by our extraction protocol, it favours the recovery of polyP. This could be due to stabilisation of polyP by cells or cell constituents, or to partial destabilisation of polyP in the ion-rich cell-free medium.

Our method of harvesting and extraction, common for ATP and polyP, allows to calculate the polyP/ATP ratio independently of any normalisation like cell density, chlorophyll concentration or dry mass measurement. Indeed, cell density measurement can become a problem in the situations when cells aggregate, for example in P and Ca deficiencies or if pH drifts too far from neutrality (29). Cell aggregation happened during the experiments of this study with the CC-4533 strain, although no harsh treatment was applied. Cell counting thus became difficult when cells formed large aggregates (> 32 cells). For some experiments (Fig. 1 & 2), we evaluated cell density by measuring optical density at 550 nm (16). However, aggregating cells stick to spectrophotometer cuvettes and these density measurements are also prone to error. Therefore, data dispersion is larger for ATP and polyP concentrations in filtration and centrifugation based extractions than for their polyP/ATP ratio. The polyP/ATP ratio, in conjunction with the ATP and polyP concentrations, can thus be used to describe the polyP physiology of cells in various physiological and stress conditions.

Filtration retrieved less polyP than direct harvesting of cells. As it is improbable that this would come from a physiological degradation of polyP due to filtration, it is probable that the 3 µm filter lets through some very small cells or cell debris that would be richer in polyP than the average of cell population. In desynchronised cultures such as ours, cell size distribution can be very wide, ranging from around 4 to 13 µm of diameter for strains used in (30). The presence of polyP-rich debris or small cells will have to be further investigated with single cell techniques.

The majority of phosphate stored as polyP in CC-4533 is found in very long chains (Fig. 7 and (31)), but the majority of chains are short. Indeed, we measure a mean length around 200 P. This means for example that for each polyP chain of 4000 P, one needs to add 20 chains of 5 P to reach the average value of 200 P, while only adding 2.5% of total P. This lets us hypothesise that our gel in Fig. 7 does not allow to visualise the majority of small chains contained in our sample. The presence of very long and very short polyP chains in our sample could reflect the different chemical forms under which polyP exist in *C. reinhardtii* cells, namely in solution or phase-separated as granules (32,33).

## Conclusion

We have devised an optimised protocol for the joint extraction of ATP and polyP from *C. reinhardtii* cells, that will greatly benefit research on cellular bioenergetics, for which this alga is a prominent model organism. The method is efficient for both metabolites, except very short chain polyP. Being based on the robust phenol-chloroform extraction, the method can readily be applied to other microorganisms, and on any physiological or stress culture conditions.

## Declarations

Ethics approval and consent to participate – Not applicable

Consent for publication – Not applicable

## Availability of data and materials

All data generated or analysed during this study are included in this published article.

## Authors’ contributions

AB conceived and supervised the project. AB and EI designed the experiments. EI performed experiments, analysed the data and wrote the manuscript. AB corrected the manuscript. EI and AB read and approved the final manuscript.

## Acknowledgements

EI and AB thank Yves Choquet and Emanuel Sanz-Luque for interesting discussions and suggestions to improve the optimised protocol, Irelka Colina for her advice on statistical analysis, and Stephan Eberhard and Benjamin Bailleul for helpful and insightful comments on the manuscript.

